# Ketamine and its metabolite 2R,6R-hydroxynorketamine promote ocular dominance plasticity and release TRKB from inhibitory control without changing perineuronal nets enwrapping parvalbumin interneurons

**DOI:** 10.1101/2022.04.06.487292

**Authors:** Cecilia Cannarozzo, Anna Rubiolo, Plinio Casarotto, Eero Castrén

## Abstract

Ketamine has been described as a fast-acting antidepressant, exerting effects in depressed patients and in preclinical models with a rapid onset of action. The typical antidepressant fluoxetine is known to induce plasticity in the adult rodent visual cortex, as assessed by a shift in ocular dominance, a classical model of brain plasticity, and a similar effect has been described for ketamine and its metabolite 2R,6R-hydroxynorketamine (R,R-HNK). Here, we demonstrate that ketamine (at 3 or 20 mg/kg) and R,R-HNK facilitated the shift in ocular dominance in monocularly deprived mice, after 3 injections, throughout the 8-days regimen. Notably, the comparison between the treatments indicates a higher effect size of R,R-HNK compared to ketamine. Treatment with ketamine or R,R-HNK failed to influence the levels of perineuronal nets (PNNs) surrounding parvalbumin-positive interneurons. However, we observed *in vitro* that both ketamine and R,R-HNK are able to disrupt the tropomyosin-related kinase B (TRKB) interaction with the protein tyrosine phosphatase sigma (PTPσ), which upon binding to PNNs dephosphorylates TRKB. These results support a model where diverse drugs promote the reinstatement of juvenile-like plasticity by directly binding TRKB and releasing it from PTPσ regulation, without necessarily affecting PNNs deposits.

## Introduction

The primary visual cortex is a well-established system to study plasticity, as this area was early on characterized with the presence of a time window of enhanced plasticity, namely a critical/sensitive period (Hubel & Wiesel, 1970). The critical period is transient and tightly regulated by the maturation of parvalbumin-positive (PV^+^) interneurons, and its closure coincides with the formation of structures of extracellular matrix called perineuronal nets (PNNs) surrounding the interneuron soma (Hensch, 2005; Pizzorusso et al., 2002).

During the critical period, deprivation of one eye in mice results in the weakening of the responses from the deprived eye and strengthening of the responses from the open eye, causing a shift in the preferential eye-driven responsiveness or ocular dominance - OD (Gordon & Stryker, 1996). Originally it was thought that OD plasticity could only occur during the critical period, however, it is now clear that juvenile-like plasticity can be restored in adulthood in various ways, such as enzymatic removal of PNNs by chondroitinase ABC (chABC), enriched environment, and dark rearing. (Fagiolini et al., 1994; Greifzu et al., 2014; Pizzorusso et al., 2002). In addition, treatment with many antidepressant compounds, including the selective-serotonin reuptake inhibitor fluoxetine (Maya Vetencourt et al., 2008; Steinzeig et al., 2017), tianeptine (Castrén & Antila, 2017), and ketamine (Casarotto et al., 2021; Grieco et al., 2020; Venturino et al., 2021) promote plasticity in the adult rat and mouse visual cortex. Notably, we have recently demonstrated that several different antidepressants, including fluoxetine, ketamine, and its metabolite 2R,6R-hydroxynorketamine (R,R-HNK) directly bind to the Brain-derived neurotrophic factor (BDNF) receptor TRKB, thereby inducing antidepressant and plasticity-like behavioral responses (Casarotto et al., 2021). Ketamine also acts as a non-competitive N-methyl-D-aspartate glutamate receptor (NMDARs) antagonist, however, many other NMDA receptor antagonists have failed to produce similar effects in clinical and preclinical trials (Abdallah et al., 2015; Autry et al., 2011; Maeng et al., 2008; Zarate, Singh, Carlson, et al., 2006; Zarate, Singh, Quiroz, et al., 2006). Moreover, blockade of NMDARs compromises plasticity in the visual cortex rather than promoting it (Kleinschmidt et al., 1987). R,R-HNK does not appear to inhibit NMDARs at the concentrations found in the brain following systemic administration (Lumsden et al., 2019; Zanos et al., 2016), but it strongly promotes a shift in ocular dominance in adult mice (Casarotto et al., 2021).

Protein tyrosine phosphatase sigma (PTPσ or PTPRS) binds to chondroitin-sulfate proteoglycans (CSPGs), major components of PNNs, and exerts a restrictive role on plasticity by dephosphorylating TRKB (Kurihara & Yamashita, 2012). PNNs are essential for the stabilization of the excitatory/inhibitory balance of the network (Lensjø et al., 2017), and their reduction by different interventions results in the reinstatement of plasticity in the visual cortex, as well as in different brain areas (Karpova et al., 2011; Pizzorusso et al., 2002; Sale et al., 2007). Interestingly, fluoxetine disrupts the interaction between PTPσ and TRKB, allowing a permissive environment for plasticity despite PNNs presence (Lesnikova et al., 2021).

Given that ketamine is only partially metabolized into R,R-HNK (Zanos et al., 2016), in this study we compared the effectiveness of these drugs in the promotion of ocular dominance plasticity. In addition, ketamine treatment has been previously associated with region-specific changes in PNNs state (Matuszko et al., 2017; Venturino et al., 2021), thus, we assessed the effects of ketamine and R,R-HNK treatment on PNNs surrounding PV^+^ interneurons in the visual cortex. Finally, considering that fluoxetine affects TRKB interaction with PTPσ and that ketamine and R,R-HNK directly interact with TRKB like fluoxetine (Casarotto et al., 2021; Lesnikova et al., 2021), we explored their effects on the interaction between TRKB and PTPσ *in vitro*.

## Methods

### Animals

For ketamine and R,R.HNK doses of 10 mg/kg, we reanalyzed the data of previously published studies (Casarotto et al., 2021). For ketamine doses of 3 and 20 mg/kg, female C57BL/6JRccHsd mice were used (Envigo-Harlan Labs, Netherlands) as in our previous studies (Casarotto et al., 2021; Steinzeig et al., 2017). The mice were group-housed (3-6 per cage - type ll: 552 cm2 floor area, Tecniplast, Italy) under a 12h light/dark cycle, with free access to water and food. All the mice were 17-20 weeks old (119-140th postnatal day, PD) at the time of the monocular deprivation, as the physiological critical period is expected to have already closed (Lehmann & Löwel, 2008). All experiments were performed in compliance with institutional guidelines and approved by the State Regional Administrative Agency for Southern Finland (ESAVI/10300/04.10.07/2016; ESAVI/38503/2019).

### Drugs

Ketamine hydrochloride (Ketalar 50mg/ml, Pfizer) at 3 or 20 mg/kg doses, diluted in sterile saline, were administered through i.p. injection three times on alternate days, during the 7-day monocular deprivation protocol (see figure 1A for timeline). 2R,6R-Hydroxynorketamine hydrochloride (R,R-HNK) was purchased from Tocris (#6094) and administered at 10mg/kg dose diluted in sterile saline. Both drugs were diluted in DMSO for the *in vitro* experiments. The doses/concentrations were chosen based on the literature (Casarotto et al., 2021; Pryazhnikov et al., 2018).

**Figure 1.**
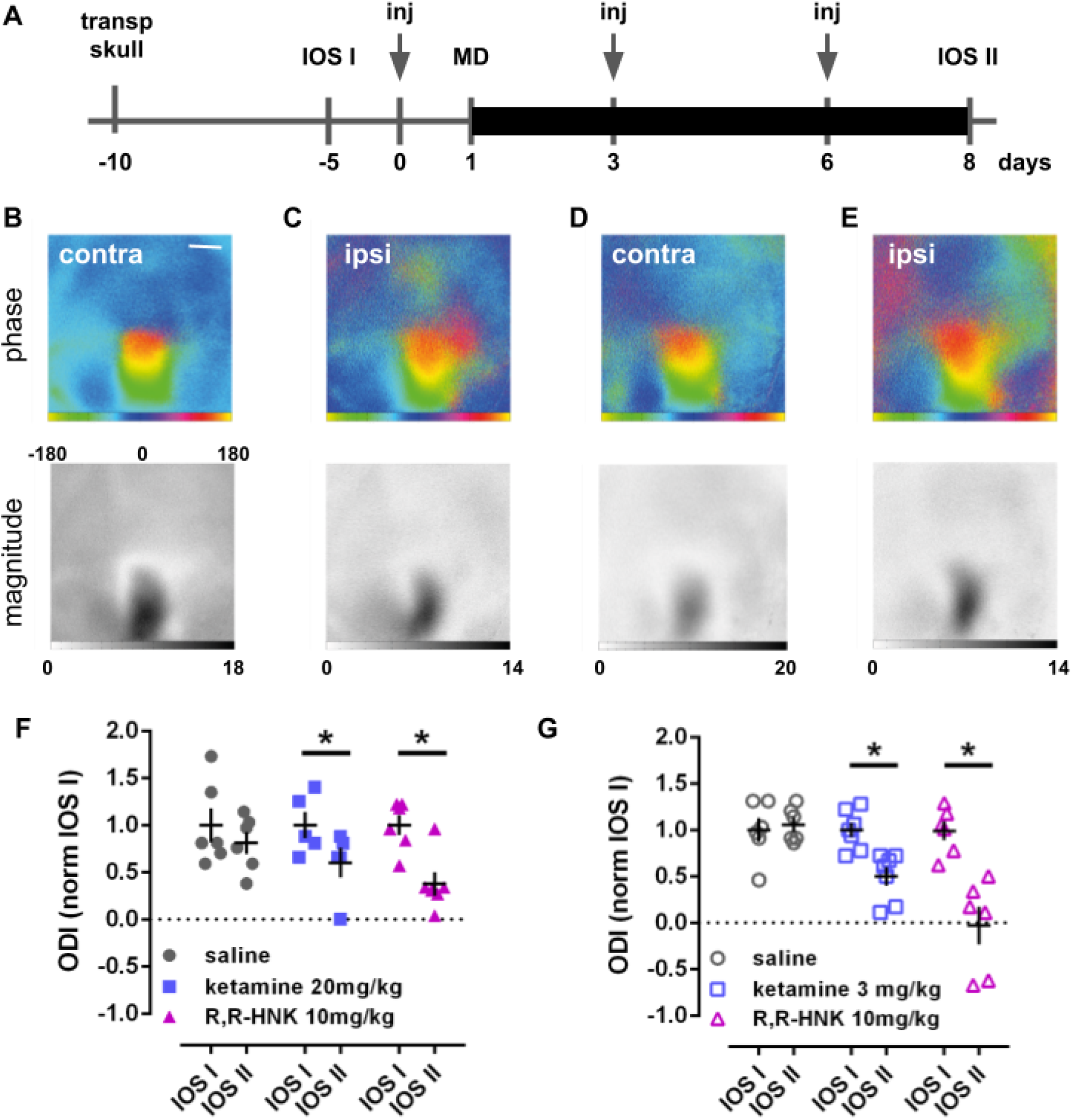
Systemic administration of ketamine in combination with monocular deprivation promotes a shift in ocular dominance index (ODI) in adult mice. **(A)** Experimental design of ketamine-induced shift in ocular dominance, thick black line indicates days under monocular deprivation (MD). **(B-E)** representative maps of intrinsic signal in the binocular region of the visual cortex from contralateral (deprived) and ipsilateral eye stimulation before (IOS-I: B, C) and after (IOS-II: D, E) ketamine (3mg/kg) treatment. Phase maps of retinotopy (upper panels), the color wedge shows the phase color coding in degrees used to visualize the response evoked by the stimulus. Magnitude maps (lower panels), the grayscale shows the amplitude of the intrinsic signal. Scale bar, 1 mm. **(F)** The administration of ketamine at 20mg/kg (yellow squares, d=1.20) promoted a shift in MD-induced ODI after 3 i.p. injections. A higher effect was observed after R,R-HNK at 10mg/kg (green triangles, d=2.20) under the same administration regimen. **(G)** The administration of ketamine at 3mg/kg (yellow squares, d=2.14) promoted a shift in MD-induced ODI after 3 i.p. injections, and a higher effect was observed after R,R-HNK at 10mg/kg (green triangles, d=2.73) under the same administration regimen. Data expressed as ocular dominance index (ODI) normalized by IOS-I. *p<0.05 from IOS-I (n=5-7/group, Fisher’s LSD).

### Surgical procedure

The transparent skull surgeries were performed as described in detail before (Steinzeig et al., 2017). Briefly, the mice were anesthetized with an anesthetic mixture containing 0.05mg/kg fentanyl (Hameln, Germany), 5mg/kg midazolam (Hameln, Germany), 0.5mg/kg medetomidine (Orion Pharma, Finland) diluted in saline (B. Braun, Germany). The eyes of the mice were protected from drying with eye-protective gel (Alcon, UK), and the mice were head-fixed in a stereotaxic frame, with a maintained temperature of 37° C. The scalp and periosteum above the visual cortex were removed and the skull was rapidly cleaned with a cotton swab soaked in acetone and air-dried. A thin layer of cyanoacrylate glue (Loctite 401, Henkel, Germany) was applied to the surface of the skull, followed by two layers of acryl, prepared by mixing colorless acrylic powder (EUBECOS, Germany) with methacrylate liquid (Densply, Germany). The mice were awakened with 0.1mg/kg buprenorphine (Indivior, UK), 1mg/kg atipamezole (VetMedic, Finland), and subcutaneously injected with 5mg/kg carprofen (VetScan) for postoperative analgesia. The day after, the mice were anesthetized with isoflurane, a metal bar holder was glued and fixed with a mixture of cyanoacrylate glue and dental cement (Densply, Germany) to the skull and transparent nail polish (#72180, Electron Microscopy Sciences) was applied above the visual cortex, visible at the center of the holder.

### Monocular deprivation

Mice were monocularly deprived for one week, anesthesia was induced by 4% isoflurane and maintained at 2% until the end of the procedure. Antibiotic gel (Isathal Vet 1%, Dechra, Canada) was applied on the contralateral eye with respect to the hemisphere to be imaged, the eyelashes were cut and the eye was sutured shut with 3 mattress sutures (Ethiconn). Antibiotic ointment (Oftan Dexa-Chlora, Anten, Finland) was applied on the sutured eye, and carprofen (5mg/kg) was injected s.c. for postoperative analgesia. All animals were checked daily and those showing signs of corneal injury were excluded from the experiments.

### Optical Imaging

Optical imaging of the intrinsic signal was performed as previously described, to determine the strength of neuronal responses to visual stimulation of either eye in the binocular visual cortex (Cang et al., 2005; Kalatsky & Stryker, 2003). Briefly, two imaging sessions were conducted: the first one before the beginning of the ketamine administration (IOS-I) followed by a second session on the 8th day after monocular deprivation (IOS-II). For imaging, anesthesia was induced with 1.8% isoflurane and then maintained with 1.2% isoflurane in a 1:2 mixture of O2/air. Animals were kept on a heating pad, 25 cm in front of the stimulus monitor. The visual stimulus, a 2° horizontal bar moving upwards at 0.125 Hz temporal frequency and 1/80 degree position frequency, was displayed in the central part of a high refresh rate monitor (width: -15° to 5° azimuth, relative to the animal visual field) in order to preferentially stimulate the binocular part of the visual field. The continuous-periodic stimulation was synchronized with a continuous frame acquisition, collected independently for each eye (Kalatsky & Stryker, 2003). First, a green light (540±20nm wavelength) was used to take vascular maps. Then, the camera was focused 600 μm below the cortical surface and red light illumination (625±10nm wavelength) was used to record the intrinsic signals for 5 minutes, at a rate of 30 frames per second. Following the acquisition of both contralateral (C) and ipsilateral (I) eyes, the cortical maps were computed after spatial binning and thresholding (figure 1B-E), and the Ocular Dominance (OD) score as (C−I)/(C+I) was calculated. The Ocular Dominance Index (ODI) was determined as the mean of the OD score for all responsive pixels. The ODI values oscillate between -1 and +1, where positive values indicate a contralateral bias, while negative ones indicate ipsilateral bias and ODI values of 0 indicate that inputs from ipsilateral and contralateral eyes are equally strong. Finally, the ODI values were normalized by values obtained in the IOS-I session.

### Sample collection

In the 20 mg/kg experiment, mice were perfused with PFA 4% immediately after the second imaging ended, whereas in the 3mg/kg experiment mice were perfused the day after. The perfused brains were post-fixed in PFA 4% overnight and then stored at 4°C in PBA.

### Immunohistochemistry

Brains were transferred to 30% sucrose solution (PBS 0,1%) for 48-72h, then included in Tissue-Tek O.C.T. compound (Sakura) and snap-frozen in dry ice. Brains were sliced coronally at the cryostat (Leica CM3050S) with a thickness of 40 µm, mounted on slides (Thermo Scientific Superfrost plus microscope slides), and stored at -20°C. Frozen sections were rinsed with PBS 1X for 10 min and blocked with 5% normal goat serum (NGS) and 0.3% Triton X-100 (Sigma) in PBS 1X for 1hr at RT. Slides were then incubated in a humidity chamber for 48h at 4°C with a primary antibody cocktail of biotin-conjugated *Wisteria floribunda agglutinin*, WFA 1:200 (Sigma Aldrich, L1516) and guinea pig anti-parvalbumin 1:500 (Synaptic System, 195 004), diluted in 1X PBS with 1% NGS and 0.3% Triton X-100. After four rounds of 10 min washings in PBS 1X, the slides were incubated in a humidity chamber for 2h at RT with a secondary antibody cocktail composed of Streptavidin Alexa fluorophore 555 conjugate 1:400 (ThermoFisher Scientific, S21381) and goat anti-guinea pig Alexa fluorophore 647 1:200 (Abcam, ab150187), in 1% NGS and 0.3%Triton X-100 (Sigma) in PBS 1X and protected from light. Slides were washed four times with PBS 1X and then incubated in a humidity chamber with Hoechst 1:10000 diluted in PBS 1X to label the nuclei and better identify the cortical layers. Finally, slides were washed with PBS 1X for 10 mins and coverslipped with Dako fluorescence mounting medium (Dako North America).

### Image acquisition and analysis

The experimenter was blind to experimental groups. The visual cortex (V1) was determined according to the Mouse brain atlas (Paxinos and Franklin, second edition) and sections in the interval from -4.36 to -3.52 from Bregma were chosen for image acquisition. A total of 3 slices containing both hemispheres were imaged with a Zeiss LSM 710 confocal microscope. 15 optical images with a Z-stack of 1 μm step were acquired per slice with a PlanApochromat 10X/0.45 M27 and imaging Software ZEN 2012 lite (Zeiss). The same microscope and camera settings were used for the imaging of all of the slides. Image processing was done using ImageJ software (National Institutes of Health, Bethesda, MA, USA). A total of 6 images per animal were counted (3 slices, both hemispheres). The region of interest (ROI) was selected in the primary visual cortex and subdivided into four layers (first, second, and third together, the fourth alone, and then fifth and sixth together). The same ROI was used to analyze all the images. The number of parvalbumin-positive (PV+), PNN-positive (PNN+), and colocalized PV+/PNN+ cells were manually counted, using the Cell Counter tool of ImageJ. Cells were considered positive and counted if they displayed a PV staining throughout the soma for the PV+, and a clear ring around the soma for the PNN+, and all the neurons fulfilling these criteria were counted as positive regardless of their size or intensity of staining. All cells entirely within the boundaries of the area were counted.

### Cell culture, sample collection and enzyme-linked immunosorbent assay (ELISA)

Cultures of hippocampal cells from rat embryos were prepared as previously described in detail (Sahu et al., 2019). Briefly, suspended hippocampal cells were seeded in poly-L-lysine-coated 24-well plates (View Plate 96, PerkinElmer) at 125000 cells/well. The cells were maintained in neurobasal medium, supplemented with B27 and left undisturbed, except for medium change (1/3 twice per week). At DIV8-10, the cells were treated with ketamine or R,R-HNK (10μM/15min), then washed with ice-cold PBS and lysis buffer was added [20mM Tris-HCl; 137mM NaCl; 10% glycerol; 0.05M NaF; 1% NP-40; 0.05mM Na3VO4], containing a cocktail of protease and phosphatase inhibitors (Sigma-Aldrich, #P2714 and #P0044, respectively).

The interaction between TRKB and phosphatase sigma (PTPσ) was determined by ELISA (Lesnikova et al., 2021). Briefly, the primary antibody against TRKB (R&D; #AF1494; 1:1000 in carbonate buffer, pH 9.8, Na2CO3 57.4mM, NaHCO3 42.6mM) was incubated in a white 96-well plate (OptiPlate 96 F HB, White, PerkinElmer) overnight (ON) at 4°C. On the second day, the plate was blocked with 2% BSA in PBS with 0.1% Tween (PBS-T) for 2h at RT. The samples were added to the plate and left ON at 4°C. The wells were washed again with PBS-T followed by incubation with secondary antibody against PTPσ (Santa Cruz, #sc-100419) ON at 4°C. Following another washing step, the plate was incubated with HRP-conjugated goat anti-mouse IgG (BioRad, #1705047) for 2h at RT. The plate was washed with PBS-T and the chemiluminescent signal was detected by a plate reader (Varioskan Flash, Thermo Scientific) after the addition of ECL (1:1). The signal from the samples, after blank subtraction, were normalized by the average of the control group and expressed as a percentage from the control.

### Statistical analysis

The OD plasticity data was obtained from independent experiments, thus the ODI from IOS-I and IOS-II sessions were submitted to paired t-test and the effect size was calculated for each of the ketamine doses used, as well as for the previously published data, using the method of Cohen’s d (Cohen, 1988). To determine a putative difference between R,R-HNK and ketamine treatments, the IOS values were normalized by IOS-I, the normalized values were analyzed by 2-way ANOVA with repeated measures, with IOS sessions and drug treatment as factors, followed by Fisher’s LSD *post hoc* test when appropriate. Stainings were analyzed with a 2-way ANOVA. P<0.05 was considered statistically significant, and all data used in the present study is stored and available in FigShare under a CC-BY license, DOI:10.6084/m9.figshare.12278606.

## Results

Our previously published data indicate that ketamine and its metabolite R,R-HNK induce an OD shift in response to monocular deprivation at a dose of 10 mg/kg (Casarotto et al., 2021). R,R-HNK produced a clear response that closely resembles that produced by chronic treatment with fluoxetine (Casarotto et al., 2021). However, the response to ketamine was more restricted than that to R,R-HNK (Casarotto et al., 2021). We re-analyzed and calculated Cohen’s d for the results of a previous study describing the effect of ketamine and R,R-HNK at 10 mg/kg in the same treatment regimen (3 i.p. injections over 8 days) in the monocular deprivation (MD)-induced shift in ocular dominance. The administration of ketamine 10 mg/kg induced a shift in ocular dominance [t(4)=6.496, p=0.003, n=5, d=2.13]. Similarly R,R-HNK also promoted a shift in ocular dominance, although with a higher effect size [t(3)=6.951, p=0.0061, n=4, d=3.22]; data from (Casarotto et al., 2021).

Since R,R-HNK represents the metabolite of only one enantiomer of the racemic ketamine, we first tested whether ketamine at a higher dose might produce the same effect as R,R-HNK did at 10 mg/kg. We therefore tested whether ketamine at 20 mg/kg might produce an OD shift that is comparable to that produced by R,R-HNK at 10 mg/kg. The drug treatment regimen is depicted in figure 1A. Following the transparent skull surgery, the animals were submitted to the first IOS session (representative images for phase and magnitude signal of contra and ipsilateral eye in figure 1B,C) and received the first injection of ketamine (20mg/kg), R,R-HNK (10mg/kg) or saline 24h before the MD procedure. The MD surgery was performed 48h after IOS-I and the eyes were maintained closed for 7 days until the end of the experiment. Two days after MD surgery the animals received a second injection, followed by another administration on day 6, 48 h before the IOS-II session (figure 1D,E). Each drug treatment was performed in independent groups, using experimentally naive mice. We observed an overall trend between treatment and trials and a significant difference between trials [interaction: F(2,14)=2.745, p=0.0987; trials: F(1,14)=26.38, p=0.0002]. The administration of ketamine 20 mg/kg induced a significant shift in ODI between IOS-I and IOS-II sessions [Fisher’s LSD, p=0.0151, n=5, d=1.20] that was similar to that produced by ketamine at 10 mg/kg, but again with a weaker effect than the effect of R,R-HNK [Fisher’s LSD, p=0.0003, n=6, d=2.20], and no difference was observed between IOS-I and IOS-II in saline-treated animals [Fisher’s LSD, p=0.177, n=6, d=0.52], figure 1F.

Then, we reasoned that even though ketamine might promote plasticity through TRKB, the action of ketamine on the NMDA receptor might counteract these effects, and therefore tested a smaller dose of ketamine (3 mg/kg). The overall analysis indicated a significant interaction between treatment and trial, as well as a significant difference between the treatments and between the trials [interaction: F(2,16)=12.87, p=0.0005; trial: F(1,16)=33.29, p<0.0001; treatment: F(2,16)=7.963, p=0.004]. Ketamine at 3mg/kg also induced a significant shift in ODI [Fisher’s LSD, p=0.0024, n=7, d=2.14], but with a smaller effect-size than R,R-HNK [Fisher’s LSD, p< 0.0001, n=6, d=2.73], and again no difference was observed between IOS-I and IOS-II in saline-treated animals [Fisher’s LSD, p=0.7103, n=6, d=-0.23], as seen in figure 1G.

We then measured the percentage of PV^+^ interneurons displaying PNNs around their soma in the visual cortices of the experimental mice. A general analysis revealed no significant difference in the percentage of PV^+^ cells surrounded by PNNs between saline, ketamine 20 mg/kg and R,R-HNK, in any of the cortical layers [2-way ANOVA, p>0.05, n=5-6], figures 2A. Similarly, no significant difference in the percentage of PV^+^ cells surrounded by PNNs was found between saline, ketamine 3 mg/kg and R,R-HNK, in any of the layers [2-way ANOVA, p>0.05, n=4-7], figure 2B.

**Figure 2.**
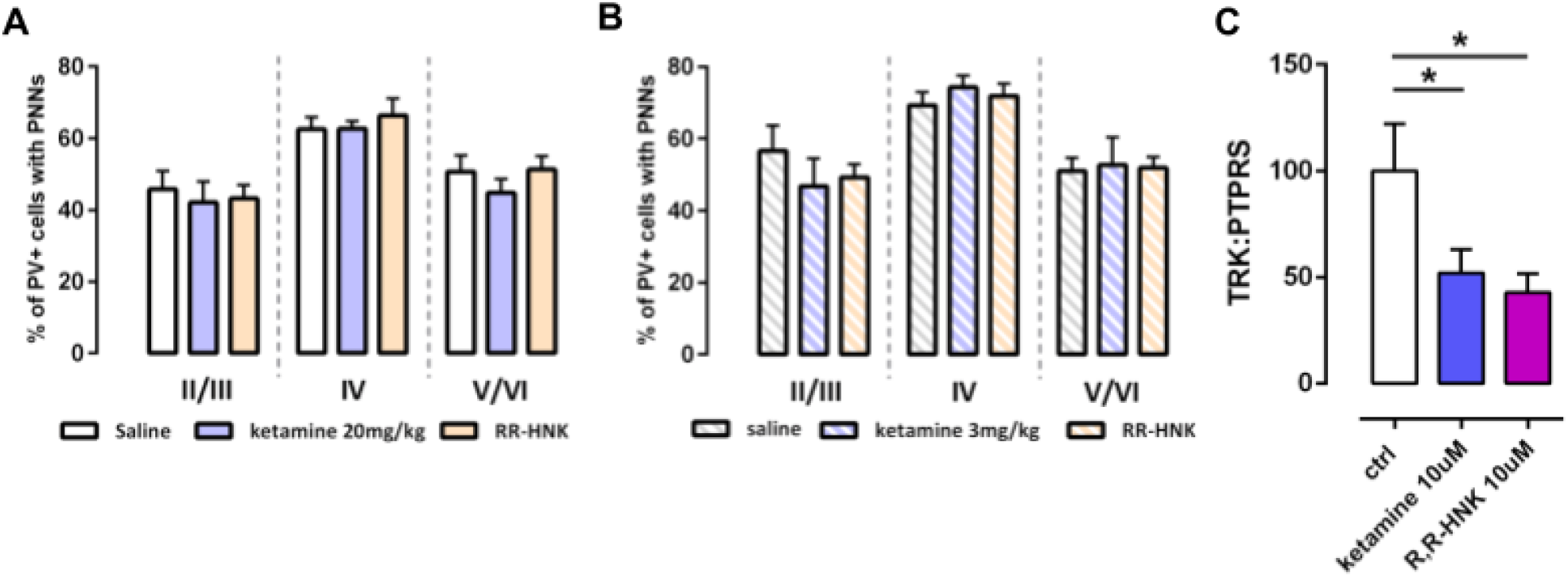
Ketamine or R,R-HNK treatment does not affect the presence of PNNs around PV^+^ cells *in vivo* but disrupts TRKB interaction with PTPσ *in vitro*. **(A)** The percentage of PV^+^ cells surrounded by PNNs did not differ in mice treated with ketamine 20 mg/kg or R,R-HNK compared to saline, in all the layers of the visual cortex. **(B)** The percentage of PV^+^ cells surrounded by PNNs did not differ in mice treated with ketamine 3 mg/kg or R,R-HNK compared to saline, in all the layers of the visual cortex. **(C)** In cultured hippocampal cells from rat embryos, the treatment with ketamine or R,R-HNK (10 μM/15min) induces disruption of TRKB:PTPσ interaction. * p<0.05 from ctrl group.

We recently demonstrated that PTPσ, a tyrosine phosphatase that acts as a receptor for components of PNN, couples to TRKB and restricts its activity, and that disruption of PNN or TRKB:PTPσ interaction is associated with reactivation of ODP (Lesnikova et al., 2021). We then tested if ketamine and R,R-HNK are able to disrupt TRKB:PTPσ interaction, as observed for fluoxetine. Our one-way ANOVA indicates a significant effect of the drug treatment on TRKB:PTPσ levels in cultured hippocampal cells [F(2,19)=3.850, p=0.0395]. The *post-hoc* analysis indicates a significant difference between ketamine or R,R-HNK compared to the vehicle-treated group (Fisher’s LSD, vehicle vs ketamine or vs R,R-HNK: p<0.05, n=7-8), figure 2C.

## Discussion

The present study demonstrates that ketamine promotes the shift in ocular dominance in adult mice and disrupts the interaction between TRKB and PTPσ, but it did not produce any measurable changes in the PNNs surrounding PV^+^ cells. These data propose a mechanism for the ketamine and R,R-HNK action that is similar to that produced by fluoxetine, where the drugs directly bind to TRKB and hinder its interaction with PTPσ, thus facilitating TRKB-dependent plasticity.

During the critical period, an MD induces a shift in ocular dominance in the primary visual cortex of mice towards the non-deprived eye (Gordon & Stryker, 1996; Hensch, 2005). After the complete closure of the critical period (>PD120 in mice), preventing the input from one eye has little to no effect on the ocular dominance (Lehmann & Löwel, 2008). Several factors can anticipate or delay the closure of the sensitive period in the visual cortex, but in general terms, this window of plasticity is associated with the maturation of GABAergic PV^+^ interneurons and the assembly of PNNs around their soma (Hensch, 2005, 2014).

Contrary to classical antidepressants that require several weeks to act, ketamine and R,R-HNK are able to induce ocular dominance plasticity (Casarotto et al., 2021; Grieco et al., 2020) and behavioral effects after a much shorter latency (Autry et al., 2011; Girgenti et al., 2017; Krystal et al., 2013; Zarate, Singh, Carlson, et al., 2006). We have recently shown that ketamine and R,R-HNK as well as fluoxetine directly bind to TRKB with a low affinity and that TRKB binding mediates the plasticity-related effects produced by antidepressants (Casarotto et al., 2021). The affinity of ketamine to TRKB is similar to its affinity to NMDARs (Casarotto et al., 2021; Zanos et al., 2018). Ketamine readily penetrates the brain to achieve concentrations needed to bind to TRKB and NMDARs, but for fluoxetine, concentrations needed to reach TRKB binding build up gradually over several days and weeks of treatment (Karson et al., 1993), which may at least partially explain the need of long-term treatment with fluoxetine to promote plasticity.

It is currently unclear which of the effects of ketamine are mediated by TRKB positive allosteric modulation and which by inhibition of NMDARs. Our recent data strongly argue that at least the plasticity-related effects, including ODP, are mediated by the direct TRKB binding. Here, we showed that different doses of ketamine produce a similar partial effect on the reopening of visual cortex plasticity. Considering the comparable affinity of ketamine to TRKB and NMDARs, and given that blockade or knock-out of NMDARs compromises visual plasticity (Fong et al., 2020; Kleinschmidt et al., 1987; Sawtell et al., 2003), our results suggest that ketamine promotes visual plasticity through TRKB, and at the same time counteracts this action through inhibition of NMDARs. This is consistent with our observation that R,R-HNK which possesses only a low affinity to NMDARs (Zanos et al., 2016), turned out to be more effective than any of the ketamine doses tested in promoting the ocular dominance plasticity (Casarotto et al., 2021).

Perineuronal nets are mesh-like structures that consist of chondroitin sulfate proteoglycans (CSPGs), hyaluronan, and tenascin-R and aggregate around soma and dendrites of a variety of neurons during late development (Brückner et al., 1993, 2006). In the visual cortex, PNNs promote the inhibitory activity of PV^+^ interneurons, participating in the regulation of the excitatory-inhibitory balance of the network and in the closure of the critical period (Lensjø et al., 2017). The reduction in PNN deposit around PV^+^ interneurons produced by enzymatic cleavage through chABC has been associated with the reinstatement of plasticity in the visual cortex, as well as other brain regions such as the amygdala, hippocampus, and frontal cortex (Gogolla et al., 2009; Pizzorusso et al., 2002). Chronic fluoxetine treatment that also reactivates critical period-like plasticity in the visual cortex has been shown to reduce the density of PNNs surrounding PV^+^ interneurons (Karpova et al., 2011; Ohira et al., 2013; Winkel et al., 2021). However, in the visual cortex, there have been also contradictory reports on the action of fluoxetine on the PNNs, which might be resulting from the different treatment protocols (Ohira et al., 2013; Steinzeig et al., 2019). New evidence indicates that various dose regimens with ketamine change the total amount of PNNs or those enwrapping PV^+^ interneurons in different brain regions, suggesting that ketamine effects on PNNs might be region-dependent and dose-dependent (Lavender et al., 2020; Matuszko et al., 2017; Venturino et al., 2021). Our results indicate that ketamine at 3 mg/kg and 20 mg/kg fails to decrease PNNs surrounding PV+ cells after 7 of MD, even though both doses successfully restore OD plasticity in the visual cortex of adult mice. In addition, we report that similarly to ketamine, the reinstatement of OD plasticity through R,R-HNK seems not to be associated with changes in PNN deposits.

We recently showed that the reactivation of OD plasticity produced by chronic fluoxetine treatment requires TRKB activity in PV+ interneurons and that direct optical activation of TRKB specifically in the PV+ neurons enables OD plasticity by downregulating the activity of PV^+^ cells, thereby promoting network activity through disinhibition (Winkel et al., 2021). We have further shown that the reactivation of OD plasticity produced by chABC-dependent PNNs disruption also relies on the activity of TRKB receptors in the PV^+^ interneurons (Lesnikova et al., 2021). Protein tyrosine phosphatase sigma (PTPσ), a ligand of CSPGs in PNNs, interacts with TRKB and compromises its activity; disruption of PNNs inhibits PTPσ and disinhibits TRKB in PV^+^ cells (Kurihara & Yamashita, 2012; Lesnikova et al., 2021). We have demonstrated that fluoxetine disrupts TRKB:PTPσ interaction, promoting TRKB activity in PV^+^ cells even when PNNs remain intact. Here we report that ketamine and R,R-HNK also produce a similar response *in vitro*, disrupting TRKB:PTPσ interaction, which further supports a common mode of action for these drugs through TRKB.

Although the exact mechanism behind the effectiveness of ketamine needs further clarification, the present study provides new evidence in support of a comprehensive model where antidepressants directly induce plasticity through TRKB in PV”interneurons, even in the absence of any macroscopic changes in PNNs structure. As the latency for the effect of ketamine in OD plasticity is significantly shorter than that observed for fluoxetine (Steinzeig et al., 2017, 2019), and this time difference agrees with the delay in the appearance of the clinical antidepressant effects, these data emphasize the role of reinstated critical period-like plasticity for the antidepressant drugs.

## Acknowledgments

the authors thank Sulo Kolehmainen, Giuliano Didio, Juzoh Umemori, and Merve Fred for their technical assistance. This study was funded by grants from the Finnish Cultural Foundation (#00210254 to CC), Päivikki and Sakari Sohlberg Foundation (to CC), European Research Council (#322742), EU Joint Programme - Neurodegenerative Disease Research (JPND) CircProt (#301225 and #643417), Sigrid Jusélius Foundation, Jane and Aatos Erkko Foundation, and the Academy of Finland (#294710 and #307416). None of the funders had a role in the data acquisition, analysis, or manuscript preparation.

## Authors’ contributions

CC, AR and PC conducted the experiments designed by CC, PC and EC; CC and PC wrote the first draft, edited by EC. EC is a shareholder of Herantis Pharma, and received lecture fees from Janssen-Cilag. All other authors declare no competing interests.

## References

Abdallah, C. G., Sanacora, G., Duman, R. S., & Krystal, J. H. (2015). Ketamine and rapid-acting antidepressants: A window into a new neurobiology for mood disorder therapeutics. Annual Review of Medicine, 66, 509–523. https://doi.org/10.1146/annurev-med-053013-062946

Autry, A. E., Adachi, M., Nosyreva, E., Na, E. S., Los, M. F., Cheng, P., Kavalali, E. T., & Monteggia, L. M. (2011). NMDA receptor blockade at rest triggers rapid behavioural antidepressant responses. Nature, 475(7354), 91–95. https://doi.org/10.1038/nature10130

Brückner, G., Brauer, K., Härtig, W., Wolff, J. R., Rickmann, M. J., Derouiche, A., Delpech, B., Girard, N., Oertel, W. H., & Reichenbach, A. (1993). Perineuronal nets provide a polyanionic, glia-associated form of microenvironment around certain neurons in many parts of the rat brain. Glia, 8(3), 183–200. https://doi.org/10.1002/glia.440080306

Brückner, G., Szeöke, S., Pavlica, S., Grosche, J., & Kacza, J. (2006). Axon initial segment ensheathed by extracellular matrix in perineuronal nets. Neuroscience, 138(2), 365–375. https://doi.org/10.1016/j.neuroscience.2005.11.068

Cang, J., Kalatsky, V. A., Löwel, S., & Stryker, M. P. (2005). Optical imaging of the intrinsic signal as a measure of cortical plasticity in the mouse. Vis. Neurosci., 22(5), 685–691.

Casarotto, P. C., Girych, M., Fred, S. M., Kovaleva, V., Moliner, R., Enkavi, G., Biojone, C., Cannarozzo, C., Sahu, M. P., Kaurinkoski, K., Brunello, C. A., Steinzeig, A., Winkel, F., Patil, S., Vestring, S., Serchov, T., Diniz, C. R. A. F., Laukkanen, L., Cardon, I., … Castrén, E. (2021). Antidepressant drugs act by directly binding to TRKB neurotrophin receptors. Cell, 184(5), 1299–1313.e19. https://doi.org/10.1016/j.cell.2021.01.034

Castrén, E., & Antila, H. (2017). Neuronal plasticity and neurotrophic factors in drug responses. Molecular Psychiatry, 22(8), 1085–1095. https://doi.org/10.1038/mp.2017.61

Cohen, J. (1988). Statistical Power Analysis for the Behavioral Sciences (2nd ed.). Routledge. https://doi.org/10.4324/9780203771587

Fong, M., Finnie, P. S., Kim, T., Thomazeau, A., Kaplan, E. S., Cooke, S. F., & Bear, M. F. (2020). Distinct Laminar Requirements for NMDA Receptors in Experience-Dependent Visual Cortical Plasticity. Cerebral Cortex, 30(4), 2555–2572. https://doi.org/10.1093/cercor/bhz260

Girgenti, M. J., Ghosal, S., LoPresto, D., Taylor, J. R., & Duman, R. S. (2017). Ketamine accelerates fear extinction via mTORC1 signaling. Neurobiology of Disease, 100, 1–8. https://doi.org/10.1016/j.nbd.2016.12.026

Gogolla, N., Caroni, P., Lüthi, A., & Herry, C. (2009). Perineuronal Nets Protect Fear Memories from Erasure. Science. https://doi.org/10.1126/science.1174146

Gordon, J. A., & Stryker, M. P. (1996). Experience-dependent plasticity of binocular responses in the primary visual cortex of the mouse. The Journal of Neuroscience: The Official Journal of the Society for Neuroscience, 16(10), 3274–3286.

Grieco, S. F., Qiao, X., Zheng, X., Liu, Y., Chen, L., Zhang, H., Yu, Z., Gavornik, J. P., Lai, C., Gandhi, S. P., Holmes, T. C., & Xu, X. (2020). Subanesthetic Ketamine Reactivates Adult Cortical Plasticity to Restore Vision from Amblyopia. Current Biology, 30(18), 3591–3603.e8. https://doi.org/10.1016/j.cub.2020.07.008

Hensch, T. K. (2005). Critical period plasticity in local cortical circuits. Nature Reviews. Neuroscience, 6(11), 877–888. https://doi.org/10.1038/nrn1787

Hensch, T. K. (2014). Bistable parvalbumin circuits pivotal for brain plasticity. Cell, 156(1–2), 17–19. https://doi.org/10.1016/j.cell.2013.12.034

Hubel, D. H., & Wiesel, T. N. (1970). The period of susceptibility to the physiological effects of unilateral eye closure in kittens. The Journal of Physiology, 206(2), 419–436.

Kalatsky, V. A., & Stryker, M. P. (2003). New paradigm for optical imaging: Temporally encoded maps of intrinsic signal. Neuron, 38(4), 529–545.

Karpova, N. N., Pickenhagen, A., Lindholm, J., Tiraboschi, E., Kulesskaya, N., Agústsdóttir, A., Antila, H., Popova, D., Akamine, Y., Bahi, A., Sullivan, R., Hen, R., Drew, L. J., & Castrén, E. (2011). Fear erasure in mice requires synergy between antidepressant drugs and extinction training. Science (New York, N.Y.), 334(6063), 1731–1734. https://doi.org/10.1126/science.1214592

Karson, C. N., Newton, J. E., Livingston, R., Jolly, J. B., Cooper, T. B., Sprigg, J., & Komoroski, R. A. (1993). Human brain fluoxetine concentrations. Journal of Neuropsychiatry and Clinical Neurosciences, 5, pp.322–322.

Kleinschmidt, A., Bear, M., & Singer, W. (1987). Blockade of “NMDA” receptors disrupts experience-dependent plasticity of kitten striate cortex. Science, 238(4825), 355–358. https://doi.org/10.1126/science.2443978

Krystal, J. H., Sanacora, G., & Duman, R. S. (2013). Rapid-Acting Glutamatergic Antidepressants: The Path to Ketamine and Beyond. Biological Psychiatry, 73(12), 1133–1141. https://doi.org/10.1016/j.biopsych.2013.03.026

Kurihara, D., & Yamashita, T. (2012). Chondroitin Sulfate Proteoglycans Down-regulate Spine Formation in Cortical Neurons by Targeting Tropomyosin-related Kinase B (TrkB) Protein. Journal of Biological Chemistry, 287(17), 13822–13828. https://doi.org/10.1074/jbc.M111.314070

Lavender, E., Hirasawa-Fujita, M., & Domino, E. F. (2020). Ketamine’s dose related multiple mechanisms of actions: Dissociative anesthetic to rapid antidepressant. Behavioural Brain Research, 390, 112631. https://doi.org/10.1016/j.bbr.2020.112631

Lehmann, K., & Löwel, S. (2008). Age-dependent ocular dominance plasticity in adult mice. PLoS One, 3(9), e3120.

Lensjø, K. K., Lepperød, M. E., Dick, G., Hafting, T., & Fyhn, M. (2017). Removal of Perineuronal Nets Unlocks Juvenile Plasticity Through Network Mechanisms of Decreased Inhibition and Increased Gamma Activity. Journal of Neuroscience, 37(5), 1269–1283. https://doi.org/10.1523/JNEUROSCI.2504-16.2016

Lesnikova, A., Casarotto, P. C., Fred, S. M., Voipio, M., Winkel, F., Steinzeig, A., Antila, H., Umemori, J., Biojone, C., & Castrén, E. (2021). Chondroitinase and Antidepressants Promote Plasticity by Releasing TRKB from Dephosphorylating Control of PTPσ in Parvalbumin Neurons. Journal of Neuroscience, 41(5), 972–980.

https://doi.org/10.1523/JNEUROSCI.2228-20.2020

Lumsden, E. W., Troppoli, T. A., Myers, S. J., Zanos, P., Aracava, Y., Kehr, J., Lovett, J., Kim, S., Wang, F.-H., Schmidt, S., Jenne, C. E., Yuan, P., Morris, P. J., Thomas, C. J., Zarate, C. A., Moaddel, R., Traynelis, S. F., Pereira, E. F. R., Thompson, S. M., … Gould, T. D. (2019). Antidepressant-relevant concentrations of the ketamine metabolite (2R,6R)-hydroxynorketamine do not block NMDA receptor function. Proceedings of the National Academy of Sciences of the United States of America, 116(11), 5160–5169. https://doi.org/10.1073/pnas.1816071116

Maeng, S., Zarate, C. A., Du, J., Schloesser, R. J., McCammon, J., Chen, G., & Manji, H. K. (2008). Cellular Mechanisms Underlying the Antidepressant Effects of Ketamine: Role of α-Amino-3-Hydroxy-5-Methylisoxazole-4-Propionic Acid Receptors. Biological Psychiatry, 63(4), 349–352. https://doi.org/10.1016/j.biopsych.2007.05.028

Matuszko, G., Curreli, S., Kaushik, R., Becker, A., & Dityatev, A. (2017). Extracellular matrix alterations in the ketamine model of schizophrenia. Neuroscience, 350, 13–22. https://doi.org/10.1016/j.neuroscience.2017.03.010

Maya Vetencourt, J. F., Sale, A., Viegi, A., Baroncelli, L., De Pasquale, R., O’Leary, O. F., Castrén, E., & Maffei, L. (2008). The antidepressant fluoxetine restores plasticity in the adult visual cortex. Science (New York, N.Y.), 320(5874), 385–388. https://doi.org/10.1126/science.1150516

Ohira, K., Takeuchi, R., Iwanaga, T., & Miyakawa, T. (2013). Chronic fluoxetine treatment reduces parvalbumin expression and perineuronal netsin gamma-aminobutyric acidergic interneurons of the frontal cortex in adultmice. Molecular Brain, 6(1), 43. https://doi.org/10.1186/1756-6606-6-43

Pizzorusso, T., Medini, P., Berardi, N., Chierzi, S., Fawcett, J. W., & Maffei, L. (2002). Reactivation of Ocular Dominance Plasticity in the Adult Visual Cortex. Science, 298(5596), 1248–1251. https://doi.org/10.1126/science.1072699

Pryazhnikov, E., Mugantseva, E., Casarotto, P., Kolikova, J., Fred, S. M., Toptunov, D., Afzalov, R., Hotulainen, P., Voikar, V., Terry-Lorenzo, R., Engel, S., Kirov, S., Castren, E., & Khiroug, L. (2018). Longitudinal two-photon imaging in somatosensory cortex of behaving mice reveals dendritic spine formation enhancement by subchronic administration of low-dose ketamine. Scientific Reports, 8(1), 1–12. https://doi.org/10.1038/s41598-018-24933-8

Sahu, M. P., Nikkilä, O., Lågas, S., Kolehmainen, S., & Castrén, E. (2019). Culturing primary neurons from rat hippocampus and cortex. Neuronal Signaling, 3(2), NS20180207. https://doi.org/10.1042/NS20180207

Sale, A., Maya Vetencourt, J. F., Medini, P., Cenni, M. C., Baroncelli, L., De Pasquale, R., & Maffei, L. (2007). Environmental enrichment in adulthood promotes amblyopia recovery through a reduction of intracortical inhibition. Nature Neuroscience, 10(6), 679–681. https://doi.org/10.1038/nn1899

Sawtell, N. B., Frenkel, M. Y., Philpot, B. D., Nakazawa, K., Tonegawa, S., & Bear, M. F. (2003). NMDA receptor-dependent ocular dominance plasticity in adult visual cortex. Neuron, 38(6), 977–985.

Steinzeig, A., Cannarozzo, C., & Castrén, E. (2019). Fluoxetine-induced plasticity in the visual cortex outlasts the duration of the naturally occurring critical period. The European Journal of Neuroscience, 50(10), 3663–3673. https://doi.org/10.1111/ejn.14512

Steinzeig, A., Molotkov, D., & Castrén, E. (2017). Chronic imaging through “transparent skull” in mice. PLoS ONE, 12(8). https://doi.org/10.1371/journal.pone.0181788

Venturino, A., Schulz, R., De Jesús-Cortés, H., Maes, M. E., Nagy, B., Reilly-Andújar, F., Colombo, G., Cubero, R. J. A., Schoot Uiterkamp, F. E., Bear, M. F., & Siegert, S. (2021). Microglia enable mature perineuronal nets disassembly upon anesthetic ketamine exposure or 60-Hz light entrainment in the healthy brain. Cell Reports, 36(1), 109313. https://doi.org/10.1016/j.celrep.2021.109313

Winkel, F., Ryazantseva, M., Voigt, M. B., Didio, G., Lilja, A., Llach Pou, M., Steinzeig, A., Harkki, J., Englund, J., Khirug, S., Rivera, C., Palva, S., Taira, T., Lauri, S. E., Umemori, J., & Castrén, E. (2021). Pharmacological and optical activation of TrkB in Parvalbumin interneurons regulate intrinsic states to orchestrate cortical plasticity. Molecular Psychiatry, 1–10. https://doi.org/10.1038/s41380-021-01211-0

Zanos, P., Moaddel, R., Morris, P. J., Georgiou, P., Fischell, J., Elmer, G. I., Alkondon, M., Yuan, P., Pribut, H. J., Singh, N. S., Dossou, K. S. S., Fang, Y., Huang, X.-P., Mayo, C. L., Wainer, I. W., Albuquerque, E. X., Thompson, S. M., Thomas, C. J., Zarate, C. A., & Gould, T. D. (2016). NMDAR inhibition-independent antidepressant actions of ketamine metabolites. Nature, 533(7604), 481–486. https://doi.org/10.1038/nature17998

Zanos, P., Moaddel, R., Morris, P. J., Riggs, L. M., Highland, J. N., Georgiou, P., Pereira, E. F. R., Albuquerque, E. X., Thomas, C. J., Zarate, C. A., & Gould, T. D. (2018). Ketamine and Ketamine Metabolite Pharmacology: Insights into Therapeutic Mechanisms. Pharmacological Reviews, 70(3), 621–660. https://doi.org/10.1124/pr.117.015198

Zarate, C. A., Singh, J. B., Carlson, P. J., Brutsche, N. E., Ameli, R., Luckenbaugh, D. A., Charney, D. S., & Manji, H. K. (2006). A randomized trial of an N-methyl-D-aspartate antagonist in treatment-resistant major depression. Archives of General Psychiatry, 63(8), 856–864. https://doi.org/10.1001/archpsyc.63.8.856

Zarate, C. A., Singh, J. B., Quiroz, J. A., De Jesus, G., Denicoff, K. K., Luckenbaugh, D. A., Manji, H. K., & Charney, D. S. (2006). A double-blind, placebo-controlled study of memantine in the treatment of major depression. The American Journal of Psychiatry, 163(1), 153–155. https://doi.org/10.1176/appi.ajp.163.1.153

